# Collagen fibre-mediated mechanical damage increases calcification of bovine pericardium for use in bioprosthetic heart valves

**DOI:** 10.1101/2020.10.21.347047

**Authors:** Alix Whelan, Elizabeth Williams, Emma Fitzpatrick, Bruce Murphy, Paul S. Gunning, David O’Reilly, Caitríona Lally

## Abstract

In cases of aortic stenosis, bioprosthetic heart valves (BHVs), with leaflets made from glutaraldehyde fixed bovine pericardium (GLBP), are often implanted to replace the native diseased valve. Widespread use of these devices, however, is restricted due to inadequate long-term durability owing specifically to premature leaflet failure. Mechanical fatigue damage and calcification remain the primary leaflet failure modes, where glutaraldehyde treatment is known to accelerate calcification. The literature in this area is limited, with some studies suggesting mechanical damage increases calcification and others that they are independent degenerative mechanisms. In this study, specimens which were non-destructively pre-sorted according to collagen fibre architecture and then uniaxially cyclically loaded until failure or 1 million cycles, were placed in an *in-vitro* calcification solution. Measurements of percentage volume calcification demonstrated that the weakest specimen group (those with fibres aligned perpendicular to the load) had statistically significantly higher volumes of calcification when compared to those with a high fatigue life. Moreover, SEM imaging revealed that ruptured and damaged fibres presented binding sites for calcium to attach; resulting in more than 4 times the volume of calcification in fractured samples when compared to those which did not fail by fatigue. To the authors’ knowledge, this study quantifies for the first time, that mechanical damage drives calcification in commercial-grade GLBP and that this calcification varies spatially according to localised levels of damage. These findings illustrate that not only is calcification potential in GLBP exacerbated by fatigue damage, but that both failure phenomena are underpinned by the unloaded collagen fibre organisation. Consequently, controlling for GLBP collagen fibre architecture in leaflets could minimise the progression of these prevalent primary failure modes in patient BHVs.

## 1. Introduction

Aortic valve stenosis (AS) remains the most prevalent heart valve disease in the Western world, and is expected to increase in the coming years due to the worldwide ageing population [1], [2]. In severe symptomatic AS patients, survival at 2 years post-diagnosis is approximately 50% [3]. AS disrupts the hemodynamics across the aortic valve and thus blood efflux from the heart with each cardiac cycle. Transcatheter aortic valve replacement (TAVR) offers minimally invasive intervention for suitable patients, by delivering a bioprosthetic heart valve (BHV) to the diseased site. These devices typically consist of a metallic frame to which three leaflets made from glutaraldehyde-fixed bovine pericardium (GLBP) are attached. Bovine pericardium is a highly collagenous tissue, where exposure to glutaraldehyde both crosslinks the collagen and sterilises the tissue; permitting its use in patient devices [4].

Although TAVR devices offer superior hemodynamics and significantly reduced intra-operative trauma in comparison to traditional mechanical valves, limited BHV durability remains a contraindication for their widespread use, especially in younger AS patients [5]. Premature device deterioration, owing specifically to the GLBP leaflets, is understood to be the primary cause of early and late BHV failure [6].

BHV leaflets exhibit two principal failure mechanisms: mechanical fatigue and calcification. Leaflets are subjected to constant cyclic loading *in-vivo*, where the loading pattern across the leaflet is multi-modal and extremely complex. We have previously reported the significant role of collagen fibre architecture on the mechanical behaviour of GLBP, where it is possible that the lack of standardised pre-screening of leaflets results in areas of inadequate fibre alignment with respect to *in-vivo* loading [7], [8]. Currently, pre-screening to ascertain the fibre patterns of GLBP is not required by ISO 5840 for BHV leaflets [8]. Moreover, it is known that glutaraldehyde accelerates the calcification of GLBP; inherently increasing its calcifying potential in patients [9], [10]. The literature in this area is both limited and contradictory where some studies propose that mechanical damage induces and accelerates tissue calcification [11]–[13]. However, other studies suggest that these are two completely independent mechanisms in BHV leaflets [14], [15]. It is also important to note that these studies have been conducted on both porcine aortic valve and bovine pericardial leaflets, which have markedly different microstructures [11]–[15].

It is understood that glutaraldehyde treatment modifies the phosphorous-rich structures in GLBP tissue, where the majority of calcification specifically occurs in devitalised cells [16]–[18]. It is also reported that free aldehyde groups, owing to the glutaraldehyde treatment, in combination with circulating calcium and phospholipids result in a passive calcification progress [19]. However, collagen and elastin fibres can also serve as binding sites for calcium, independent of cellular structures [20]–[23]. Interestingly, increased calcification is associated with younger patients. Although the correlation between age and calcium deposition is well-known, the underlying cause for this is not [17]. It has also been observed in clinically explanted BHVs that regions of high mechanical stress in the leaflets are associated with increased levels of calcification, yet the mechanism underpinning this relationship is not currently known [13], [24]. Calcification of leaflets can result in significantly sub-optimal performance of BHVs in patients. Clusters of calcium deposits result in highly localised stress concentrations, while calcification in general reduces the mobility and flexibility of the leaflets moving from systole to diastole [6], [25]. The inability of GLBP leaflets to fully open increases the systolic gradient across the valve, thereby reducing valve effective orifice area (EOA) and thus the volume of blood ejected from the heart. For diastole, the inability to fully coapt results in regurgitation, causing valvular insufficiency. Amongst others, these occurrences are classified clinically as evidence of a degenerative and dysfunctional BHV [6].

To investigate how mechanical damage influences the calcification of commercial-grade GLBP, dogbone specimens, which were previously uniaxially loaded to failure or 1 million cycles, were placed in an *in-vitro* calcifying solution and the resulting percentage volume of calcium was measured. Each specimen was non-destructively pre-sorted according to collagen fibre orientation and dispersion using small angle light scattering (SALS), allowing for a correlation between collagen fibre architecture, mechanical damage and calcification progression. As we have demonstrated that the mechanical response, and thus damage accumulation, of GLBP is dictated by the organisation of collagen fibres, here we investigate if the collagen fibre-mediated mechanical damage influences GLBP’s propensity for calcification [7], [26]. Finally, the calcified tissue samples were imaged under microscopy to visually ascertain the distribution of calcium across damaged samples.

## 2. Methods

### 2.1. Sample preparation

Dogbone specimens were cut from commercial-grade GLBP tissue that was obtained from Boston Scientific Corporation (Galway, Ireland). Each dogbone was non-destructively imaged using a small angle light scattering (SALS) system and grouped according to collagen fibre orientation and alignment as outlined previously in an earlier study [7]. Briefly, each tissue specimen was imaged at a 250 μm step-size, across a 6 x 1 mm region of analysis, centred on the dogbone gauge length. A series of images (n = 96) were analysed in Matlab (The Mathworks, MA, USA), where only specimens meeting pre-defined fibre orientation and alignment criteria proceeded to mechanical testing. Specimen categories are as follows; XD (collagen fibres highly aligned perpendicular to the uniaxial load), HD (collagen fibres highly dispersed in many directions, i.e. no dominant fibre angle) or PD (collagen fibres highly aligned parallel to the uniaxial load). Table 1 below summarises the collagen fibre angle and alignment threshold criteria for each group, where alignment is measured as the eccentricity of the elliptical scattered light distributions of each recorded image.

**Table 1:** Fibre angle and alignment criteria for XD, HD and PD groups. Note: angle is given with respect to the uniaxial load direction. Due to a very low alignment value for HD specimens, the measured fibre angle is not applicable here.

| Group | Fibre angle ($^{\circ}$) | Alignment |
| --- | --- | --- |
| XD | $90 \pm 15$ | $> 0.70$ |
| HD | N/A | $< 0.65$ |
| PD | $0 \pm 15$ | $> 0.7$ |

### 2.2. Uniaxial cyclic tensile tests

A total of 21 (n = 7 for each group) specimens which met the fibre architecture specifications (see Table 1) were mechanically tested under uniaxial cyclic tensile conditions. In a Tytron Microforce System (MTS Systems Corporation, MN, USA), each specimen was loaded to a peak stress of 3 MPa (R-ratio = 0.1), until failure or 1 million cycles was reached, at 1.5 Hz. Testing was conducted in a saline bath at 37°C to simulate physiological conditions. A supraphysiological peak stress of 3 MPa was chosen to accumulate mechanical damage in a shorter timeframe. This is also conducted in ‘Dynamic Mode Failure Testing’ as per ISO 5840, where the transvalvular pressure is increased to induce failure earlier in accelerated wear testing (AWT) [8]. 3 MPa places the stress-strain response of GLBP in the collagen-dominant region, where lower loads result in a tissue-matrix dominant response (initial linear portion of the stress-strain curve; see [7]). Thus, a stress of 3 MPa permits the accumulation of collagen fibre damage in this study, and is lower than the mean ultimate tensile strength of the weakest specimen category, namely XD [7].

A detailed analysis of the results from the aforementioned mechanical testing is found in [26]. Briefly, all PD specimens reached one million cycles without failure, and four of seven HD specimens also reached 1 million cycles without failure. In contrast, XD specimens had a statistically significantly lower fatigue performance; four of seven specimens failed after one loading cycle (see Figure 1, p < 0.0001). There was no statistically significant difference in the fatigue behaviour between HD and PD groups.

**Figure 1:**
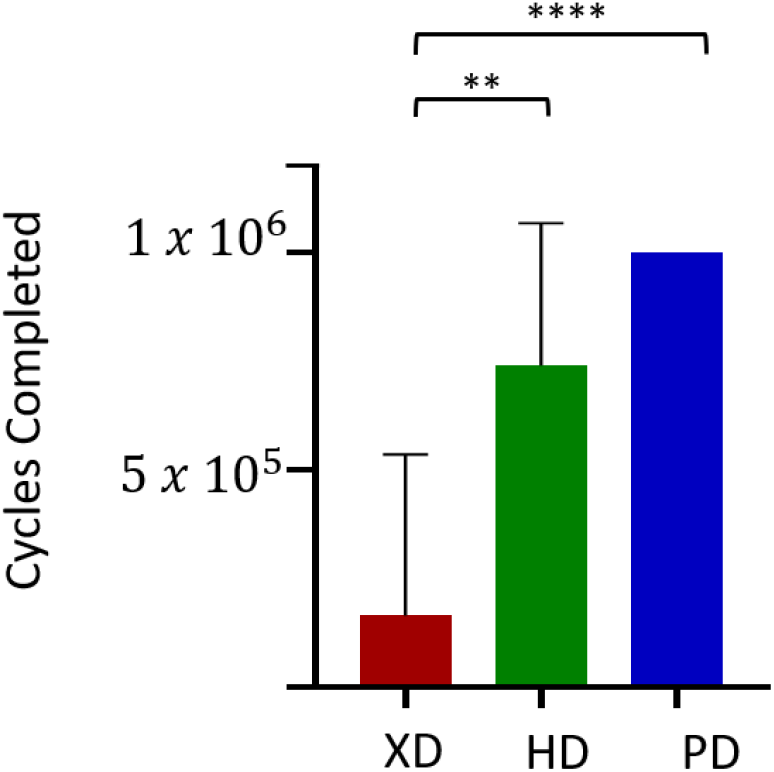
Cycles completed for XD, HD and PD specimen groups (n=7), **p <0.01, ****p <0.0001. Adapted from [26].

### 2.3. In-vitro calcification & quantification

#### 2.3.1. Calcification treatment

Following uniaxial cyclic tensile loading, each specimen from our previous study was placed in an *in-vitro* calcification solution [27]. Additionally, seven (n=7) new unloaded samples of highly dispersed (HD) and highly aligned (HA; i.e. XD/PD) fibre architecture were also placed in the calcification solution to serve as undamaged controls. This calcium-phosphate treatment calcifies specimens over 42-days, where the solution is replaced every 7 days. The concentrations of calcium and phosphate in the incubation solution (prepared in 0.05 M Tris buffer, pH = 7.4) were 3.87 mM and 2.32 mM respectively, yielding a ratio of Ca/P04 = 1.67, as in hydroxyapatite [27]. Each specimen was placed in a calcification-solution filled vial and agitated by rotation in a 37°C bath.

Each specimen was rinsed in PBS after calcification treatment, to remove excess surface calcium deposits, and measured for percentage volume calcium content within 72 hours.

#### 2.3.2. Calcification quantification

Calcium content was measured non-destructively on the gauge length of each dogbone specimen, using Micro Computed Tomography (MicroCT40, Scanco Medical, Brüttisellen, Switzerland). MicroCT uses X-ray imaging in 3D at a very high resolution, allowing for the quantification of calcium in the bulk samples. MicroCT has previously been used to measure calcification in pericardial tissue [28]. Imaging was performed at a voltage of 70 kV, and a current of 113 μA, where images were taken across approximately 200 slices per sample. With a threshold of 90, sigma of 0.8 and support of 1, 3D reconstruction and calcium quantification were achieved. In this study, MicroCT results are presented as calcification volume, as a percentage of total tissue volume. A one-way ANOVA was used to investigate any statistically significant differences between the percentage volume calcification of XD, HD and PD groups (n=7 for each group, 95 % confidence interval).

To validate the MicroCT set-up and parameters, a calcium assay (Sentinel Calcium Kit 17667, Sentinel Diagnostics, Milan, Italy) was performed on twenty (n = 20) specimens which had previously been analysed with MicroCT. Each tissue specimen was cut into smaller sections, frozen in liquid nitrogen and ground using a mortar and pestle. Following this the samples were digested in a 1M HCL solution at 60°C and 10 rpm. Once digested, the samples were assessed using the calcium assay in accordance with the manual. Briefly, the concentration of calcium was quantified based on the principle that Cresolphtalein Complexone (CPC) reacts with calcium ions at a pH < 10 to form a red colour complex. The colour intensity is directly proportional to the concentration of calcium in the sample and was measured using a plate reader set to 570 nm. All samples (calcium standard controls and tissues specimens) were assessed in triplicate and the calcium concentration was extrapolated from the averaged readings entered into a standard curve. To compare the results from the MicroCT to the calcium assay, a linear regression analysis was completed at a 95% confidence interval.

#### 2.3.3. Tissue Imaging

Representative specimens from testing groups (fatigue-damaged and non-calcified, fatigue-damaged and calcified) were imaged via Scanning Electron Microscopy (SEM) and Helium Ion Microscopy (HIM) techniques. This allowed for visualisation of both the collagen fibres and the attached calcification. For SEM imaging, specimens were first rinsed with ultra-pure water to release residual salts from PBS storage, and then snap-frozen in liquid nitrogen at 196°C for two minutes. Next, the samples were lyophilised in a VirTis™ freeze-drier (SP Scientific, Gardiner, NY, USA), with primary drying at −10°C for 10 hours and secondary drying at 25°C for 2 hours. Each specimen was then mounted on a carbon platform for SEM analysis.

HIM imaging (Carl Zeiss Helium Ion Microscope, Carl Zeiss, Oberkochen, Germany) was conducted by mounting the tissue specimens on the carbon platform as for SEM analysis. A 22° tilt angle, 10 μm aperture, 30kV voltage and spot size of 4 were used.

## 3. Results

### 3.1. Calcification

It was observed that the weakest specimen group accumulated statistically significantly higher levels of calcification, as measured by microCT, following the 42-day *in-vitro* treatment when compared to all remaining groups (XD; see Figure 2 (a)). Specifically, the mean percentage volume calcium of the XD group was 20.47 ± 9.7% tissue volume, in contrast to 4.49 ± 5.53 % and 1.58 ± 2.99 % for HD and PD groups, respectively. Although the XD group was statistically significantly different to both the HD and PD groups, there was no significant difference between the HD and PD groups.

**Figure 2:**
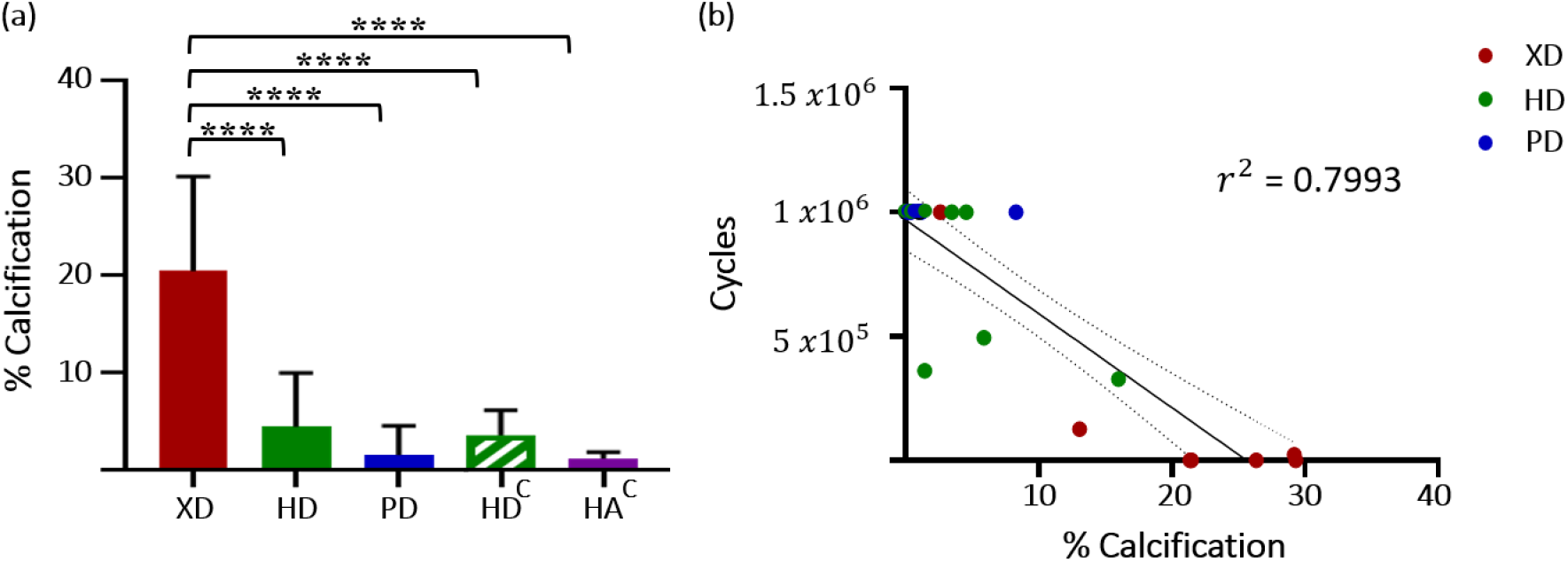
(a) % Volume calcification for XD, HD, PD, control HD and control highly aligned groups; ****p < 0.0001. (b) Linear regression analysis for cycles completed and percentage volume calcification, dotted lines indicate 95% confidence interval of regression, r^2^ = 0.7993.

Additionally, there was no statistically significant difference between HD/PD specimens and their respective undamaged controls. Moreover, the four specimens with the highest level of calcification across all twenty-one specimens failed after one loading cycle (from the XD group; see Figure 2 (b)).

Regression analysis (Figure 2 (b)) indicates a high correlation between loading cycles and calcification; that of an inverse relationship. Interestingly, the samples which reached the test end-point without failure, had levels of calcification which were not statistically significantly different to their unloaded and thus, undamaged controls.

Figure 3 (a) below shows MicroCT analysis of an unloaded control GLBP sample, where calcification is evident in dark/grey regions. Although levels of calcification are low here, it appears to be predominantly located on the tissue’s edges. In contrast, Figure 3 (b) shows MicroCT analysis of a XD specimen which failed after one loading cycle. It is clear here that the calcification is concentrated at the fracture site (black arrow), where it becomes more dispersed moving laterally to the dogbone extremities (left and right).

**Figure 3:**
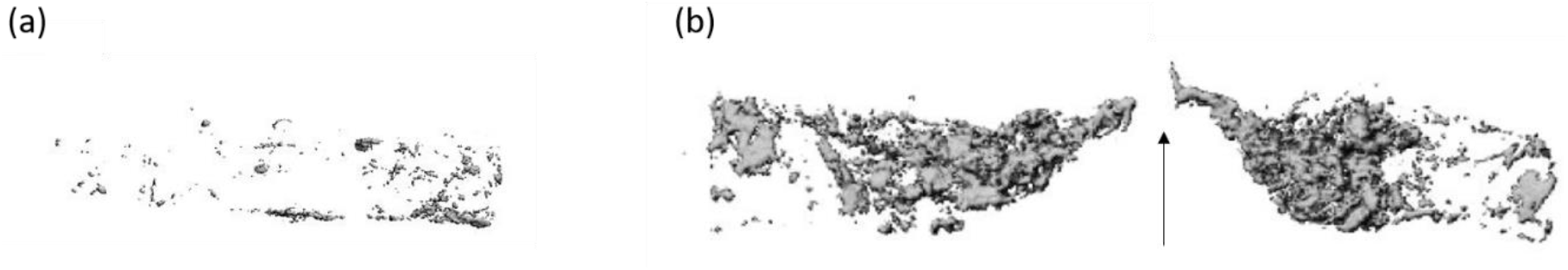
(a) MicroCT analysis of unloaded (i.e. undamaged) GLBP specimen gauge length; where darker regions indicate detected calcium, (b) XD specimen which failed from fatigue loading, where arrow indicates fracture site on gauge length centre.

Figure 4 illustrates the correlation between the MicroCT measurements and that of the Sentinel Calcium assay (*r*^2^ = 0.89); confirming the MicroCT set-up employed in this study as an accurate non-destructive alternative to the standard assay techniques.

**Figure 4:**
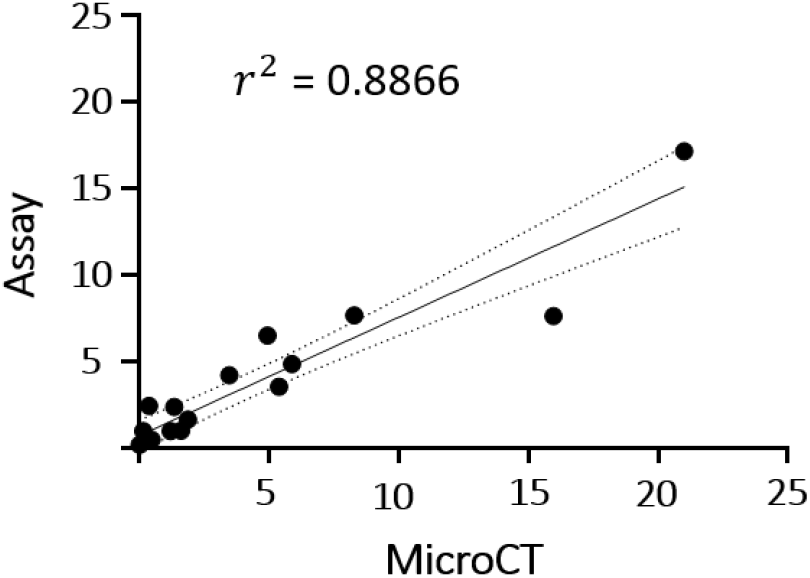
Regression analysis of % volume calcification measurements from Sentinel Calcium assay and MicroCT techniques (see section 2.3.2, r^2^ = 0.89, dotted lines indicate 95% confidence interval)

### 3.2. Tissue Imaging

Figure 5 (a) shows a specimen from the PD group after fatigue loading (1 million cycles), showing the uniformly aligned fibres along the gauge length, in agreement with the fibre orientation and alignment as measured previously through SALS. Figure 5 (c,d) shows another PD sample after fatigue loading and calcification treatment; where the calcium is clearly visible in the bright/white regions. These images also demonstrate that calcium is specifically attached to collagen fibre bundles in the form of a coating.

**Figure 5:**
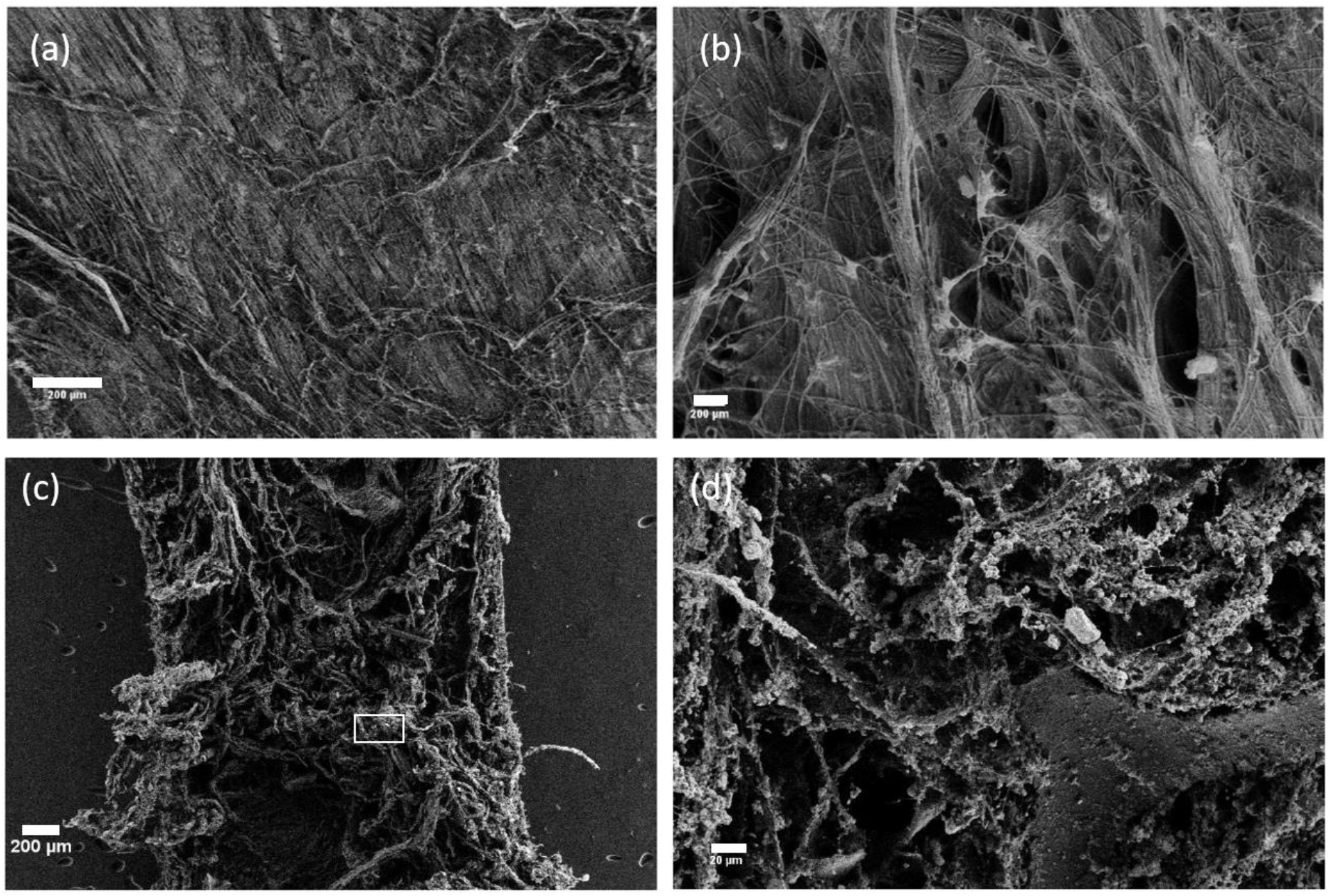
(a, b) Gauge length of uncalcified PD sample after 1 million loading cycles (scale bars both 200 μm), (c, d) gauge length of PD sample after 1 million cycles and calcification treatment, where white box in (c) indicates image location of (d), (scale bars 200 μm and 20 μm, respectively).

Figure 6 (a,b) shows an uncalcified XD sample which failed after 1 loading cycle with torn and ruptured fibres at the fracture site on the gauge length centre. Figure 6(c,d) shows another XD specimen after failure (1 cycle) and calcification treatment, with high levels of calcification (white regions). As in the PD specimen (Figure 5 (c,d)), the calcification appears to coat fibre bundles, but here we also see nodules and high concentrations attached to the tips of broken and fractured fibres (Figure 6 (d)).

**Figure 6:**
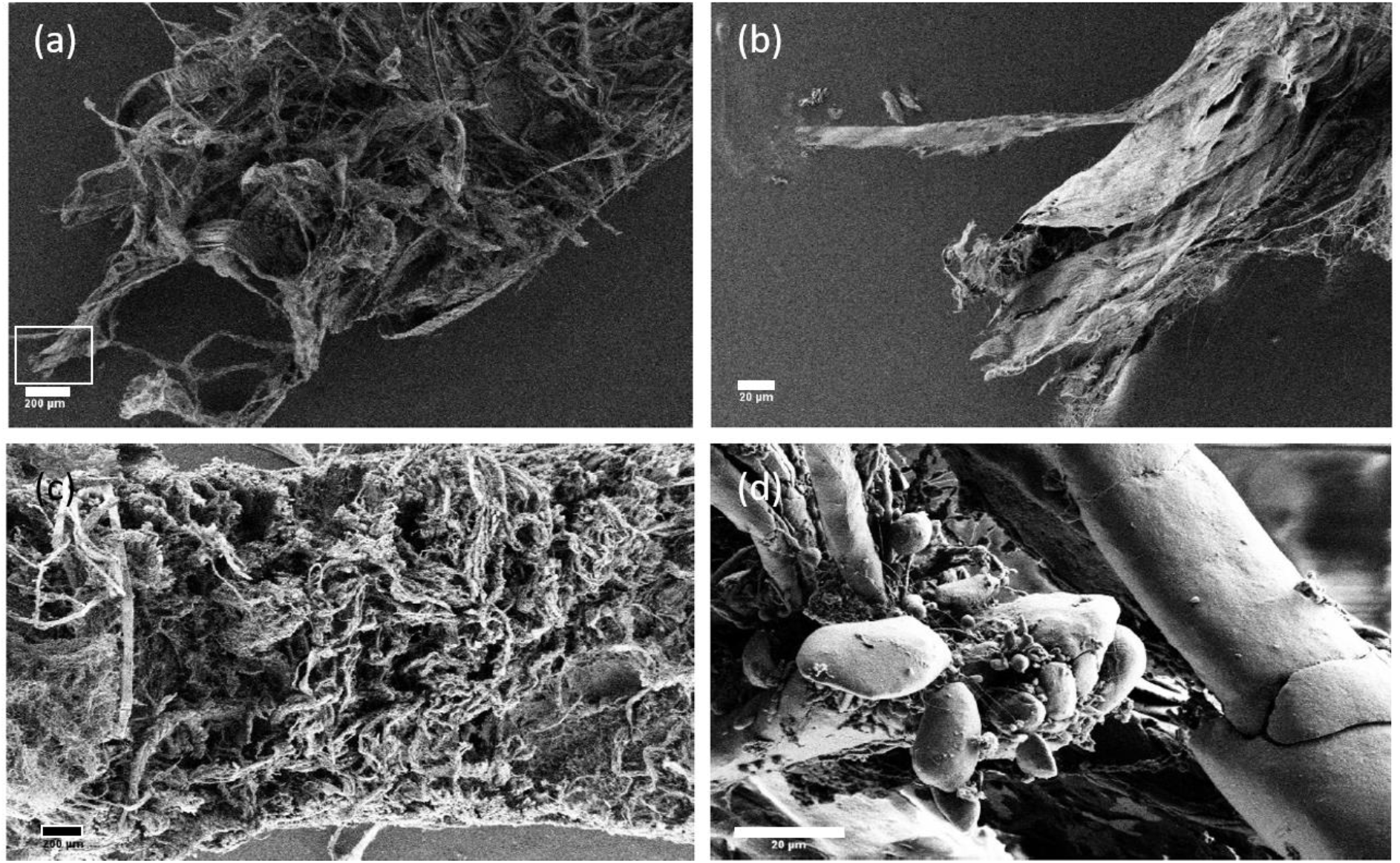
(a) Fracture site of failed, un-calcified XD sample, white box indicates image location in (b), (scale bars 200 μm and 20 μm, respectively), (c,d) failed XD sample after calcification treatment (scale bar 200 μm), where (d) shows a magnified view of attached calcium nodules (scale bar 20 μm).

Figure 7 shows representative SEM images of XD, HD and PD specimens (a-c, respectively). Comparison of these three groups indicates that calcium is attached to all regions of the XD tissue, with little contrast between raised surface fibre bundles and the more homogenous tissue below. Additionally, there appears to be more disorganised surface fibre bundles in this failed XD sample when compared to its HD/PD counterparts (Figure 7 (a-c)). Both HD and PD specimens have increased levels of calcification on surface fibres in comparison to the smooth and organised tissue fibres below.

**Figure 7:**
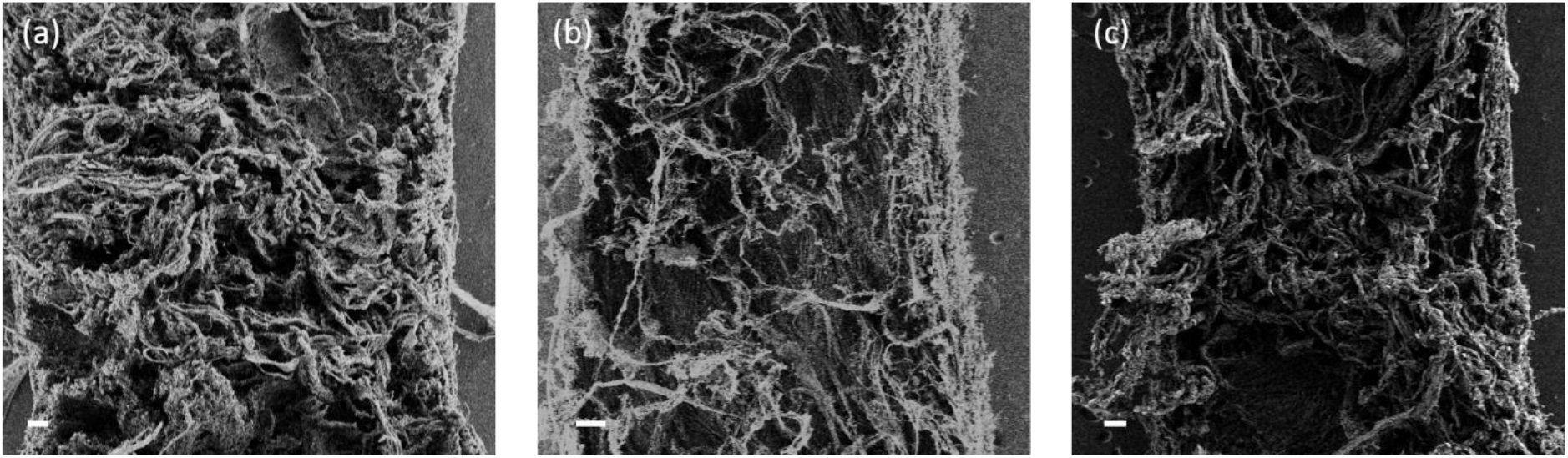
Representative (a) XD (failed after 1 cycle), (b), HD (did not fail up to 1 million cycles) and (c) PD (did not fail up to 1 million cycles) specimens after fatigue loading and calcification (all scale bars 100 μm).

Figure 8 (a, b) shows SEM analysis of an XD sample which failed after 126,006 cycles, with localised calcification bound to the extremity of a ruptured fibre bundle as also seen in Figure 6 (b). HIM analysis of the same sample allows for visualisation at an increased resolution, with calcium present in the hairpin bend of a particularly exposed fibre at the rupture site (Figure 8 (c,d)).

**Figure 8:**
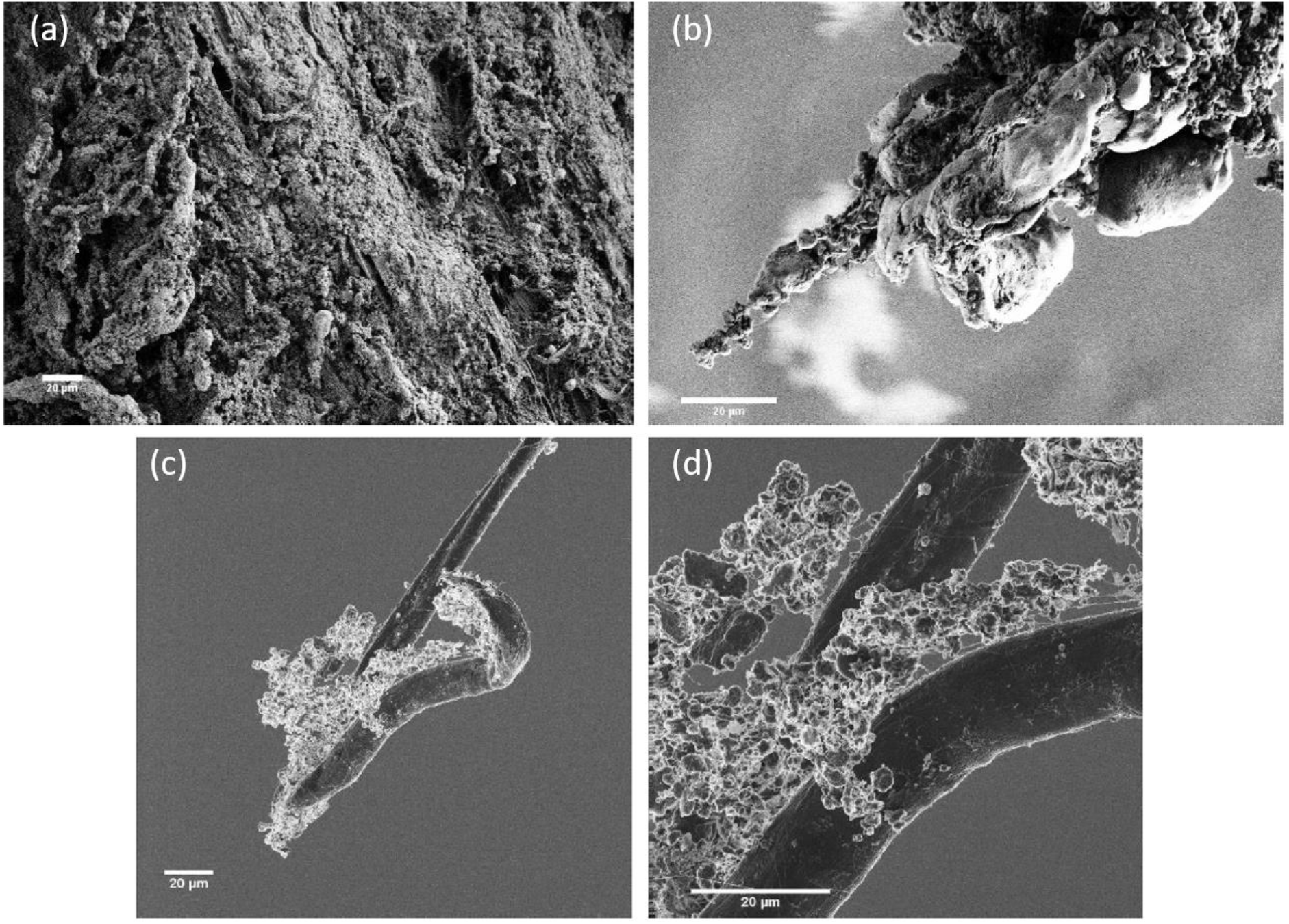
(a) SEM of failed XD specimen after calcification treatment (scale bar 20 μm), (b) magnified view of (a) showing surface fibre with calcium nodules attached (scale bar 20 μm), (c,d) HIM images of sample in (a,b), showing calcium attachment on individual fibre (scale bars both 20 μm).

## 4. Discussion

Mechanical damage and calcification are widely considered to be the primary causes of premature leaflet failure in BHVs. To improve the long-term performance of valve prostheses, it is imperative to understand the mechanisms of these failure phenomena, in addition to their relationship.

Firstly, the MicroCT and calcium assay correlate with a high r^2^ value of 0.89, indicating that MicroCT is a viable non-destructive alternative technique for measuring calcium content in GLBP samples (see Figure 4). It is not possible to non-destructively measure calcification as a function of time or its spatial organisation using traditional tissue solubilising assay techniques. However, the MicroCT techniques employed here can achieve this. This is advantageous for future experimental studies, where calcification can be measured and monitored non-destructively at incremental time points throughout a long-term experiment. Additionally, MicroCT analysis can quantify the spatial distribution of calcification on a given sample. Figure 3(b) shows that the calcification was concentrated at the fracture site on a broken GLBP specimen; i.e. the region of maximum fibre disruption and damage. This was in agreement with the SEM and HIM analysis conducted; where at much increased resolution, visualisation of calcium clusters and nodules attached to broken fibre bundles was possible (see Figure 8 (b-d)).

Figure 2 (b) demonstrates the direct relationship between mechanical damage and calcification, where load-induced damage accelerated GLBP calcification. Specifically, the XD group which completed statistically significantly less cycles than either the HD or PD groups, had a statistically significantly greater volume of calcification (see Figure 2 (a), p < 0.0001). This finding conclusively demonstrates that mechanical damage in commercial-grade GLBP accelerates calcification accumulation. Interestingly, there was no statistically significant difference between HD/PD groups, or their respective, un-damaged controls. Although not statistically significant, Figure 2 (a) shows that the mean percentage volume calcification of the unloaded HD group was greater than the unloaded highly aligned group (HA). This is possibly explained by an increased collagen content in HD GLBP; resulting in a greater number of aldehyde bonds present, and thus the potential for calcification, as calcium ions are understood to bond with aldehyde groups [26].

It has been observed in previous studies that areas of high stress, likely indicative of tissue damage, are associated with higher levels of calcification, but there remains a lack of understanding as to why this is the case [11], [13], [24]. The findings presented in this study agree with these previous studies, but the addition of calcification visualisation via MicroCT and SEM in this work elucidates the mechanism that underlies this direct relationship. Firstly, MicroCT imaging showed that calcium was predominantly located at the fracture site of a broken sample, when compared to an unloaded control GLBP sample (Figure 3). SEM imaging of both uncalcified and calcified specimens revealed that calcium adhered to the fibre bundles, where SEM analysis of undamaged and damaged GLBP indicates that broken and frayed collagen fibres offer binding sites for calcium; with increased concentrations at ruptured, exposed fibres (Figure 6 (c,d)). High stresses in these regions resulted in significant damage, providing a greater number of attachment sites and elevating the global volume of calcification to statistically significantly higher levels when compared to specimens which did not accumulate significant mechanical damage and fail (Figure 2). As described previously, glutaraldehyde treatment will inherently cause calcification of bovine pericardium, but this is exacerbated in the presence of mechanically damaged collagen fibres. Thus, the absence of significant fibre damage in specimens which did not fail by fatigue loading (i.e. HD/PD groups) limits the number of calcium binding sites. Consequently, this resulted in no statistically significant difference between HD/PD groups and their respective undamaged controls as seen in Figure 2 (a).

A limitation of this work is the uniaxial-tensile loading environment employed to induce tissue damage as this is not representative of the multi-modal and complex leaflet loading environment in-vivo. Yet, pre-sorting of the GLBP specimens according to fibre orientation and alignment with SALS allowed for direct correlation of fibre architecture and mechanical performance; and in turn how this influences calcification potential. Furthermore, as this study was conducted on commercial-grade GLBP tissue, the findings are markedly relevant to implantable devices, where commercial-grade GLBP is significantly stiffer than the GLBP tissue typically employed in the literature [7], [29].

Translating these experimental findings to GLBP leaflets presents important implications for BHV devices. A lack of fibre pre-screening in the manufacturing stage may result in collagen fibre-load malalignment in-vivo. In turn, these potentially weak areas will result in damaged fibres, thereby accelerating calcification and inducing premature valve failure as reported in the literature [6], [30], [31]. Localised regions of calcium nodules have been shown to reduce the overall mobility and flexibility of the leaflet during its transition from systole to diastole [32]. It is also important to consider the crimping of BHVs prior to delivery. It has been shown that crimping and subsequent BHV balloon inflation or self-expansion can cause leaflet injury; by disrupting the collagen fibres [33], [34]. This process may further accelerate calcification of BHV leaflets in-vivo. Furthermore, the current requirement to assess BHVs up to 200 million cycles under AWT conditions determines only the presence and accumulation of mechanical damage, but not that of calcification [8]. Perhaps AWT studies should also include assessment of calcification, thereby investigating both primary failure modes of BHVs.

Compromising GLBP leaflet functionality will have serious short and long-term consequences for BHV device viability and patient prognosis. Ultimately, this study provides fundamental insights into the relationship between the two most prevalent degenerative processes within BHVs, and demonstrates that both are underpinned by the unloaded collagen fibre architecture.

## 5. Conclusion

Mechanically induced damage was found to directly increase percentage volume calcium in commercial-grade GLBP in this study. Furthermore, we have demonstrated that collagen fibre architecture and content underpin the mechanical response and thus damage behaviour in this tissue. Premature pericardial leaflet degeneration owing to fatigue damage and calcification are the primary failure modes of BHV leaflets, yet there is currently no requirement to pre-screen for the mechanically dominant collagen fibre orientation in ISO 5840. The results presented in this study, in addition to previous work, indicate that collagen fibre architecture must be controlled for in the manufacture of BHV leaflets in order to minimise the progression of both fatigue damage and calcification *in-vivo*.

## Acknowledgements

The authors would like to thank Dermot Daly for his assistance with both the SEM and HIM imaging, carried out at the Advanced Microscopy Laboratory (AML) at the SFI centre, AMBER, CRANN Institute. This research is funded by the Irish Research Council, Boston Scientific Corporation (EBPPG/2016/353) and the European Research Council (ERC) under the European Union’s Horizon 2020 research innovation programme (Grant Agreement No. 637674).

## References

[1] V. T. Nkomo, J. M. Gardin, T. N. Skelton, J. S. Gottdiener, C. G. Scott, and M. Enriquez-Sarano, “Burden of valvular heart diseases: a population-based study,” Lancet, vol. 368, no. 9540, pp. 1005–1011, Sep. 2006.

[2] R. L. J. Osnabrugge et al., “Aortic Stenosis in the Elderly,” J. Am. Coll. Cardiol., vol. 62, no. 11, pp. 1002–1012, Sep. 2013.

[3] C. M. Otto and B. Prendergast, “Aortic-valve stenosis - From patients at risk to severe valve obstruction,” New England Journal of Medicine, vol. 371, no. 8. Massachussetts Medical Society, pp. 744–756, 2014.

[4] G. Arcidiacono, A. Corvi, and T. Severi, “Functional analysis of bioprosthetic heart valves,” J. Biomech., vol. 38, no. 7, pp. 1483–1490, Jul. 2005.

[5] H. Baumgartner et al., “2017 ESC/EACTS Guidelines for the management of valvular heart disease,” Eur. Heart J., vol. 38, no. 36, pp. 2739–2791, Sep. 2017.

[6] D. Dvir et al., “Standardized Definition of Structural Valve Degeneration for Surgical and Transcatheter Bioprosthetic Aortic Valves,” Circulation, vol. 137, no. 4, pp. 388–399, 2018.

[7] A. Whelan et al., “Collagen fibre orientation and dispersion govern ultimate tensile strength, stiffness and the fatigue performance of bovine pericardium,” J. Mech. Behav. Biomed. Mater., vol. 90, no. August 2018, pp. 54–60, 2019.

[8] International Standards Organisation, “ISO 5840-3:2013 Cardiovascular implants - cardiac valve prostheses. Part 3: Heart valve substitutes implanted by transcatheter techniques,” vol. 2013, p. 104p., 2013.

[9] G. Golomb, F. J. Schoen, M. S. Smith, J. Linden, M. Dixon, and R. J. Levy, “The role of glutaraldehyde-induced cross-links in calcification of bovine pericardium used in cardiac valve bioprostheses.,” Am. J. Pathol., vol. 127, no. 1, pp. 122–30, Apr. 1987.

[10] M. Grabenwöger et al., “Impact of glutaraldehyde on calcification of pericardial bioprosthetic heart valve material.,” Ann. Thorac. Surg., vol. 62, no. 3, pp. 772–7, Sep. 1996.

[11] M. J. Thubrikar, J. D. Deck, J. Aouad, and S. P. Nolan, “Role of mechanical stress in calcification of aortic bioprosthetic valves.,” J. Thorac. Cardiovasc. Surg., vol. 86, no. 1, pp. 115–25, Jul. 1983.

[12] K. Liao, E. Seifter, D. Hoffman, E. L. Yellin, and R. W. M. Frater, “Bovine Pericardium Versus Porcine Aortic Valve: Comparison of Tissue Biological Properties as Prosthetic Valves,” Artif. Organs, vol. 16, no. 4, pp. 361–365, Nov. 2008.

[13] G. M. Bernacca, A. C. Fisher, ’ R Wilkinson, T. G. Mackay, and D. J. Wheatley’, “Calcification and stress distribution in bovine pericardial heart valves.”

[14] M. S. Sacks and F. J. Schoen, “Collagen fiber disruption occurs independent of calcification in clinically explanted bioprosthetic heart valves,” J. Biomed. Mater. Res., vol. 62, no. 3, pp. 359–371, 2002.

[15] I. Vesely, J. E. Barber, and N. B. Ratliff, “Tissue damage and calcification may be independent mechanisms of bioprosthetic heart valve failure.,” J. Heart Valve Dis., vol. 10, no. 4, pp. 471–7, Jul. 2001.

[16] J. M. Dunn and L. M. Marmon, “Mechanisms of Calcification of Tissue Valves,” Cardiol. Clin., vol. 3, no. 3, pp. 385–396, Aug. 1985.

[17] F. J. Schoen and R. J. Levy, “Calcification of tissue heart valve substitutes: Progress toward understanding and prevention,” Ann. Thorac. Surg., vol. 79, no. 3, pp. 1072–1080, 2005.

[18] M. Valente, U. Bortolotti, and G. Thiene, “Ultrastructural substrates of dystrophic calcification in porcine bioprosthetic valve failure,” Am. J. Pathol., vol. 119, no. 1, pp. 12–21, 1985.

[19] W. Chen, F. J. Schoen, and R. J. Levy, “Mechanism of efficacy of 2-amino oleic acid for inhibition of calcification of glutaraldehyde-pretreated porcine bioprosthetic heart valves.,” Circulation, vol. 90, no. 1, pp. 323–9, Jul. 1994.

[20] R. J. Levy, F. J. Schoen, J. T. Levy, A. C. Nelson, S. L. Howard, and L. J. Oshry, “Biologic determinants of dystrophic calcification and osteocalcin deposition in glutaraldehyde-preserved porcine aortic valve leaflets implanted subcutaneously in rats.,” Am. J. Pathol., vol. 113, no. 2, pp. 143–55, 1983.

[21] F. J. Schoen, R. J. Levy, A. C. Nelson, W. F. Bernhard, A. Nashef, and M. Hawley, “Onset and progression of experimental bioprosthetic heart valve calcification.,” Lab. Invest., vol. 52, no. 5, pp. 523–32, May 1985.

[22] F. J. Schoen, J. W. Isao, and R. J. Levy, “Calcification of Bovine Pericardium Used in Cardiac Valve Bioprostheses Implicationsfor the Mechanisms ofBioprosthetic Tissue Mineralization.”

[23] M. T. Bailey, S. Pillarisetti, H. Xiao, and N. R. Vyavahare, “Role of elastin in pathologic calcification of xenograft heart valves,” J. Biomed. Mater. Res. - Part A, vol. 66, no. 1, pp. 93–102, Jul. 2003.

[24] K. K. Liao et al., “Mechanical Stress: An Independent Determinant of Early Bioprosthetic Calcification in Humans,” 2008.

[25] F. Sturla et al., “Impact of different aortic valve calcification patterns on the outcome of transcatheter aortic valve implantation: A finite element study,” J. Biomech., vol. 49, no. 12, pp. 2520–2530, Aug. 2016.

[26] A. Whelan et al., “Bovine Pericardium of High Fibre Dispersion Has High Fatigue Life and Increased Collagen Content; Potentially an Untapped Source of Heart Valve Leaflet Tissue,” Ann. Biomed. Eng., Oct. 2020.

[27] G. Golomb and D. Wagner, “Development of a new in vitro model for studying implantable polyurethane calcification,” Biomaterials, vol. 12, no. 4, pp. 397–405, 1991.

[28] A. Parekh, “Calcification Of Bovine Pericardial Aortic Heart Valves,” Electron. Thesis Diss. Repos., Aug. 2015.

[29] W. Sun, M. Sacks, G. Fulchiero, J. Lovekamp, N. Vyavahare, and M. Scott, “Response of heterograft heart valve biomaterials to moderate cyclic loading,” J. Biomed. Mater. Res., vol. 69A, no. 4, pp. 658–669, Jun. 2004.

[30] A. Kataruka and C. M. Otto, “Valve durability after transcatheter aortic valve implantation,” J. Thorac. Dis., vol. 10, no. I, pp. S3629–S3636, 2018.

[31] R. M. Reul, M. K. Ramchandani, and M. J. Reardon, “Transcatheter Aortic Valve-in-Valve Procedure in Patients with Bioprosthetic Structural Valve Deterioration,” Methodist DeBakey cardiovascular journal, vol. 13, no. 3. Methodist DeBakey Heart & Vascular Center, pp. 132–141, 01-Jul-2017.

[32] E. Jorge-Herrero, J. M. García Páez, and J. L. Del Castillo-Olivares Ramos, “Tissue heart valve mineralization: Review of calcification mechanisms and strategies for prevention,” J. Appl. Biomater. Biomech., vol. 3, no. 2, pp. 67–82, 2005.

[33] R. Zegdi, P. Bruneval, D. Blanchard, and J. N. Fabiani, “Evidence of leaflet injury during percutaneous aortic valve deployment,” Eur. J. Cardio-thoracic Surg., vol. 40, no. 1, pp. 257–259, Jul. 2011.

[34] B. Amahzoune, P. Bruneval, B. Allam, A. Lafont, J.-N. Fabiani, and R. Zegdi, “Traumatic leaflet injury during the use of percutaneous valves: a comparative study of balloon-and self-expandable valved stents,” 2012.

